# A stem cell secretome delays functional decline and supresses inflammation in two distinct models of neurodegeneration

**DOI:** 10.64898/2026.03.18.712006

**Authors:** Stuart Dickens, Andrew Parnell, Daniel Feist, Ben Mellows, Ketan Patel, Steve Ray, Samantha McLean, Robert Mitchell, Ritchie Williamson

**Affiliations:** School of Pharmacy, Optometry, and Medical Sciences, University of Bradford, United Kingdom; Micregen Ltd. Thames Valley Science Park, Reading, United Kingdom; School of Biological Sciences, University of Reading, Reading, United Kingdom

**Author notes:** Corresponding Author: Ritchie Williamson.

**Keywords:** Motor Neurone Disease, Alzheimer’s disease, Stem Cell Secretome

## Abstract

Alzheimer’s disease (AD) is a progressive neurodegenerative disorder with a rapidly increasing global prevalence. Current pharmacological interventions offer symptomatic relief but do not modify disease progression. Secretome-based therapeutics have emerged as a potential disease-modifying strategy, given their capacity to influence multiple pathological pathways, including amyloid burden, reactive gliosis, and neuronal survival. Early clinical studies support the safety and potential efficacy of these approaches, indicating mechanisms involving neuroprotection, neurodegeneration, and modulation of neuroinflammation, processes central to AD pathology. In the present study, we investigated the therapeutic efficacy of multipotent stromal cell (MSC)-derived secretomes produced by a specific platform (Secretomix) in two distinct mouse models of neurodegenerative disease: An AD model characterized by amyloid pathology, and a motor neurone disease (MND) model exhibiting TDP-43 protein aggregation. Administration of the MSC secretome resulted in a positive modulation of the behavioural phenotype in the AD model, and reduction in the rate of decline of motor co-ordination (attenuated the progression of motor deficits) in the MND model. In the latter, these functional benefits were accompanied by a measurable reduction in neuroinflammatory responses but without direct alteration of standard neuropathological markers. Additionally, ex vivo assays using human peripheral blood demonstrated broad anti-inflammatory activity of the MSC secretome, providing a potential mechanistic basis for the in vivo observations. Collectively, these findings support further investigation of MSC-derived secretomes as a promising therapeutic approach for neurodegenerative disorders, with relevance across proteinopathies characterised by distinct molecular pathways.

**Significance Statement:** Here we demonstrate the efficacy of a stem cell secretome in ameliorating cognitive and behavioural phenotypes in different models of neurodegeneration. These models represent distinct neuropathological features that are unaffected by stem cell secretome treatment but share common features of modulation of inflammation post stem cell secretome treatment. This study highlights the therapeutic potential of stem cell secretomes in the treatment of neurodegenerative conditions with an already existing neuropathology.

## 1. Introduction

Dementia represents an ongoing clinical, social, and economic challenge with no clinical options at any stage of the disease that can stop the degenerative process. It is progressively debilitating across multiple modalities (memory, thinking, behaviour and movement) and is ultimately fatal. It affects 50 million individuals worldwide in 2021 with a new diagnosis every 3 seconds (**World Alzheimer Report 2025**). Therapeutic options for Alzheimer’s disease (AD) approved across the FDA, EMA and MHRA include cholinesterase inhibitors and a glutamate receptor antagonist (**Lanctot et al, 2003; Muir et al, 2006**). These only offer symptomatic relief without addressing underlying neuropathology (**Huang et al, 2020**). Recent advances have been made in addressing underlying pathology with the approval of lecanemab and aducanumab (subsequently discontinued). While we wait validation of these interventions, development of disease targeted therapies and symptomatic therapies continues. There are currently 182 ongoing clinical trials studying the effects of novel drugs and their ability to address Alzheimer’s disease (**Cummings et al., 2025**).

Different neurodegenerative conditions are defined by their neuropathological hallmarks and clinical phenotypes. However, the boundaries between the different dementias are not always distinct, to the extent that multiple brain pathologies usually co-exist in the same individuals (**Rabinovici et al, 2016**). As such, there is a need for therapeutics that target multiple pathologies and clinical phenotypes. Establishment of pathology in the brain can occur up to 30 years before onset of symptoms (**Beason-Held et al, 2013**) and so potential therapies need to address an already existing pathology and/or the sequalae of biochemical and behavioural effects that arise during the progression of neurodegenerative disorders.

Multipotent stromal cells (MSCs) have been investigated for their potential use in regenerative medicine due to their multipotency and ability to migrate to injury sites and secrete regenerative factors (**Zhidu et al, 2024**). They have been isolated from multiple tissues, including bone marrow, adipose tissue and amniotic fluid (**Chen et al, 2008; Slamecka et al, 2016**). It is now widely accepted that the primary mechanism behind their regenerative potential is through the factors secreted by them (the secretome). This includes extracellular vesicles, nucleic acids and free protein that are able to exert beneficial effects on other cells in a paracrine manner (**Mitchell et al, 2019; Mellows et al, 2017**). Due to the diverse number of components identified within the MSC secretome, they have been investigated as a potential therapeutic strategy for a wide range of disorders, including musculoskeletal, nervous system, cardiovascular and immune-related conditions (**Maldonado et al, 2023**). This therapeutic versatility has been largely attributed to the multifaceted regenerative mechanisms of action exhibited by these components, which encompasses angiogenic, mitogenic, immunomodulatory and neurogenic functions (**Mitchell et al, 2019; Teixeira et al, 2016**).

The number of clinical trials involving MSCs has more than tripled in the last 10 years, from 493 in June 2015 to 1,582 recorded on 10^th^ July 2025. Of these, 67 were linked to three common neurological conditions: 21 in Alzheimer’s Disease (AD), 15 in Parkinson’s Disease and 31 in motor neurone disease (MND) (**Squillaro et al, 2016, clinicaltrials.gov**). Clinical trials in individuals with MND with whole MSC therapeutic candidates showed no negative side effects and improved functional rating (**Petrou et al., 2016**) and another trial reported an increase in survival time (**Barczewska et al, 2020**). Extensive literature emphasises a fundamental role for multipotent stromal cells (MSCs) in the repair and regeneration of damaged tissue, mediated through the secretome.

In this study, we investigated the therapeutic potential of MSC secretomes in 2 mouse models of neurodegenerative diseases that are characterized by accumulation of aggregates of different proteins in different brain regions: AD (amyloid pathology) and MND (TDP-43 pathology). Each model presented with only a single brain pathology representative of the disease, has a well characterised temporal cognitive and/or motor and pathological phenotype and were investigated at an age coincident with already established pathology and behavioural deficits. Primary endpoints were behavioural (mobility and/or cognition) and secondary endpoints were pathological (brain pathology/inflammation).

We report that a MSC secretome generated using the Secretomix® platform methodology from a GMP-validated amniotic fluid cell line exhibited positive modulation of the behavioural phenotype in an Alzheimer’s disease model (MRG1061), while a separate secretome produced via the same platform and cell line attenuated the rate of motor coordination decline and reduced brain inflammation in a motor neurone disease mouse model (MRG1062). Cytokine analysis revealed sex dependent alterations in mediators of inflammation in both the MSC secretome-treated Alzheimer’s mouse and motor neurone disease mouse models. Ex vivo experiments in human blood indicate that the MSC secretome has broad anti-inflammatory effects, which may underly our observed modulation of behavioural phenotypes. These results support further exploration of MSC secretome as a potential therapeutic for neurodegenerative disorders.

## 2. Materials and Methods

### 2.1 Animals

5XFAD mice overexpressing both mutant human amyloid beta (A4) precursor protein 695 (APP) with the Swedish (K670N, M671L), Florida (I716V), and London (V717I) Familial Alzheimer’s Disease (FAD) mutations and human PS1 harbouring two FAD mutations, M146L and L286V (**Oblak et al, 2021**) were purchased from Jackson Laboratories and used to test the efficacy of MRG1061. TDP-43^Q331K^ mice expressing mutant Q331K (**Mitchell et al, 2015**) were used to test efficacy of MRG1062. Animals were housed in groups of 5 in standard laboratory conditions with free access to food and water. All experimental procedures were carried out in accordance with the Animals (Scientific Procedures) Act, UK (1986) and were approved by the University of Bradford ethical review process.

### 2.2 Western Blotting

Protein extraction and Western blotting were performed as previously described (**Muha et al, 2019**). Mouse brains were rapidly dissected and frozen in liquid nitrogen and stored at –80°C. Tissue was lysed in 50 mM Tris-HCl pH 7.4, 0.1 mM EGTA, 1 mM EDTA, 1% Triton-X100, 1 mM sodium orthovanadate, 50 mM sodium fluoride, 5 mM sodium pyrophosphate, 0.27 M sucrose, 0.1% 2-mercaptoethanol supplemented with protease inhibitors (1 mM benzamidine, 0.2 mM PMSF and 5 µM leupeptin). Proteins were separated on precast 10% gels by SDS-PAGE and transferred to Nitrocellulose membrane and blocked for 1 hour at room temperature in 5% bovine serum albumin in TBST (Tris-buffered saline with 0.1% Tween-20). After blocking, membranes were incubated with primary antibody in blocking buffer overnight at 4°C with shaking. Primary antibodies were detected by incubation with R680/800-labelled secondary antibodies at room temperature for 1 h and imaged/quantified using the Li-Cor Odyssey infrared imaging system (Li-Cor) and Image Studio Lite software. The linear range of all antibodies was determined for optimal binding and concentration.

### 2.3 Secretome formulation and administration

Second trimester human amniotic fluid-derived mesenchymal stem cells (AFSC) were expanded in culture in DMEM supplemented with 10% FBS and 1% L-glutamine at 37°C, 5% CO_2_, and maintained at <70% confluence. The MRG1061 and MRG1062 formulations were generated using the Secretomix® platform methodology, proprietary approaches for producing optimised MSC secretome protocols. Briefly, cells were enzymatically harvested at confluence, washed to remove residual complete media and resuspended in saline for each secretome generation. Once each incubation process was complete, separately, each secretome was centrifuged at 300 rcf for 5 minutes to remove cells and at 2,000 rcf to remove cellular debris. The final material was sterile filtered through a 0.22 μm filter before being aliquoted for storage at –80 °C. Secretome was administered by tail vein injection every two weeks.

### 2.4 In vitro inflammation assay

RPMI media (Gibco 11875-093) was mixed with freshly collected (within 2 h) whole human blood collected at a ratio of 3:1 and 800 μL of the RPMI/blood mixture added to each well of a 24 well plate. 100 μL of secretome or vehicle was added to the respective wells and the plates incubated for 1 h at room temperature. 100 μL of 10 ng/mL LPS made up in RPMI media was added to all wells (with RPMI media alone used for unstimulated controls) and plate incubated for 5 h at 37 °C, 5% CO_2_.

Following incubation, supernatant was collected into a 1.5 mL microfuge tubes and centrifuged at 2,000 rcf for 20 min. The top layer was collected, taking care not to disturb the buffy coat or RBC layer and frozen overnight. To test individual protein concentrations, samples were tested via ELISA using R&D Systems human TNF α kit (DuoSet kit #DY210). Briefly, protein standards were prepared between 15.6 – 1,000 pg/mL and 100 μL standard/samples added to a 96 well plate and incubated for 2 h at room temperature. Concentrations were determined by Streptavidin-HRP absorbance at 540 nm.

Samples for multiplex analysis were normalised to 5 mg/mL protein with 1x PBS and sent to Eve Technologies Inc. for analysis on their Mouse Cytokine/Chemokine 32-Plex Discovery Assay® Array (MD32), 32 different beads/capture antibodies were added to each sample. Following washing of the beads to remove any unbound sample, biotinylated detection antibodies were added and incubated allowing binding to captured target molecules. Once any unbound detection antibodies are washed away, streptavidin conjugated with phycoerythrin were added. The beads were then passed through a Bio-Plex 200 analyser, which operates with a dual laser system. Combined with a standard curve, the concentration of each target molecule was calculated.

### 2.5 Behavioural testing

All animals were tested at baseline on the behavioural paradigms before undergoing treatment with secretome or saline (vehicle control). Novel object recognition (NOR) was carried out in mice (n=15/group). The number of animals per group was determined using power calculations based on results from our previous behavioural studies (**McLean et al, 2016; McLean et al, 2021; Yun et al, 2022**). Novel object recognition (NOR) was conducted separately for each individual mouse, and the location of the novel object was randomly assigned. Mice were habituated to the NOR arena (52 cm wide × 40 cm high × 52 cm long) for 20 min for three consecutive days prior to testing day. Testing day consisted of a 3 min habituation session with the mouse placed in the NOR arena followed by an acquisition trial in which two identical objects were placed in opposite corners with the animal for a period of 3 min. The mouse was then removed and placed back in the home cage for 60 min inter-trial interval (ITI). The arena was cleaned with 70% ethanol and the mouse was re-introduced to the arena for the retention trial for 3 min with a novel object and an unused triplicate version of the familiar object (previously seen in the acquisition trial).

All experiments were video recorded for subsequent analysis of behaviour. The exploration time (s) of each object in each phase was recorded manually using two stopwatches and the discrimination index was calculated: discrimination index (DI) = (time exploring the novel object(s) – time exploring the familiar object) / total time exploring both novel and familiar objects. The DI represents the difference in exploration time expressed as a proportion of the total time spent exploring the two objects in the retention trial. Locomotor activity was quantified by scoring the total number of lines crossed by the mouse during both acquisition and retention trials.

For the open field test, arena construction and set up is as described previously (**McLean et al, 2009**).

Motor coordination and strength were assessed on the rotarod apparatus (Harvard Bioscience) as described previously (**Mitchell et al, 2015**). Briefly, animals were tested at baseline, 1 month and 2 months using a 5-min acceleration protocol (4-40 rpm). Mice were tested multiple times and underwent an acclimatisation phase at 4 rpm (2 days prior to test day), followed by a training phase of steady speed (4 rpm) then an acceleration protocol (4-20 rpm over 5 min).

### 2.6 Immunohistochemistry

Brains were dissected and fixed in 4% PFA for 24 hours at 4 °C. Tissue was then dehydrated using a tissue processor (Leica) in increasing concentrations of ethanol (70-100%) and xylene before saturation with paraffin wax (Paraplast). Brains were embedded in paraffin and 5 µm sections created using the cryostat and mounted on gelatine coated slides.

The paraffin was removed by immersing the slides in xylene followed by decreasing concentrations of ethanol (100%– 70%) before washing in distilled water. Sections then underwent antigen retrieval prior to staining. Iba1 ( Proteintech #10904-1-AP), GFAP (Dako #Z0334) and p62 (Abcam #ab91526): 10mM Tris, pH 9.0 90 °C. Amyloid-β (6E10, Biolegend #803001): 30 min in 50 mM Citraconic, pH 6.0, 80 °C. Heating was performed using a water bath and sections were cooled in the buffer for 30 min then washed with water. Endogenous peroxides were quenched by submersion in 3% MeOH for 15 min then washed for 5 min in water. Slides were blocked for 60 minutes in 10% normal goat serum in TBS-Tx (0.1% Triton x-100). Slides were incubated with primary antibodies overnight at 4 °C: GFAP (1:1,000), Iba1 (1:1,000), p62 (1:1,000), 6E10 (1:5,000). Slides were washed with 3x 5 min washes in TBS-Tx then incubated with the appropriate secondary antibodies (R&D Systems VisUCyte HRP Polymer Antibody rabbit #VC001-025 or mouse #VC003-025) for 1 hour at room temperature. The wash step was then repeated and the slides developed using a DAB+ kit (Agilent #K346711-2) following the manufacturer’s instructions. Slides were then washed in distilled water then counterstained with Cresyl Violet (Fisher Scientific #0501481). Slides then washed in distilled water and dehydrated in ethanol and xylene as before, then mounted in DPX (Merck #06522).

Slides were imaged using the Motic SlideScanner and tile scanned images were analysed using QuPath.

## 3. Results

### 3.1 Effects on Behaviour

To test anxiety and locomotion in the model prior to treatment (baseline), 5XFAD mice were tested in the open field arena. In the open field test, there were no observed differences between the test group (MRG1061) and the control group (saline) in locomotion or anxiety in any of the animals tested, in that all animals recorded similar times in central squares, edge squares and corner squares suggesting that this cohort of animals all behaved similarly (Figure 1. A). The effect of MRG1061 on working memory was then investigated using the elevated Y-Maze apparatus. There was no difference between the test group (MRG1061) and the control group (saline) in spatial memory as determined by alternation rate and crossings at baseline and 1 month (data not shown). When analysed by gender, there was no difference in behaviour between male and female mice (Figure 1. A).

**Figure 1.**
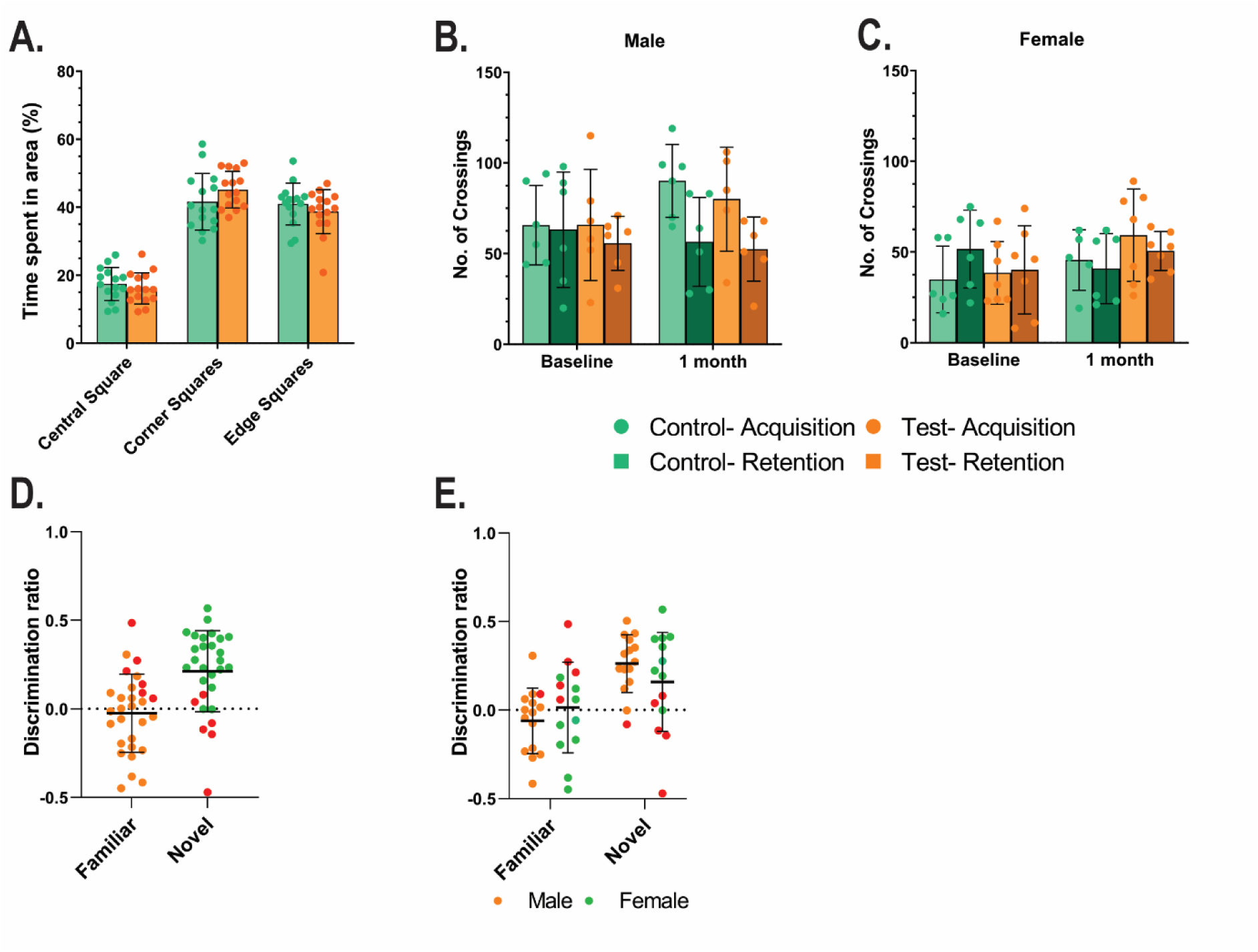
Locomotion and behaviour in 5XFAD mice. (A) Quantitation of open filed data at baseline for control (saline-treated) and (test (MRG1061-treated) groups. Open field for male and female mice. (B, C) Quantitation of locomotion of mice in NOR testing of control and test mice at baseline and one month parsed by sex. (D) Discrimination ratio of all mice at baseline. (E) Discrimination ratio at baseline of mice parsed by sec. *n*=15 male, *n*=15 female.

We then assessed mouse behaviour using the novel object recognition (NOR) test. There was no difference in locomotion between the control group and the test group at either baseline or 1 month treatment with MRG1061(Figure 1. B, C). Baseline testing revealed that while most mice could discriminate between the novel object and the familiar object (Figure 1. D), more females showed no preference or a preference for the familiar object (Figure 1. E).

As we determined that there were sex differences at baseline in preference for familiar over novel objects in the mice, we next analysed the data based on sex. There were no significant differences between MRG1061 treated and saline treated mice in exploration time of the novel or familiar object at either baseline or 1 month post treatment (Figure 1. A, B, E, D). Analysis of the discrimination index revealed that MRG1061-treated male mice could still discriminate between a novel object and a familiar object after 1 month whereas saline-treated male mice displayed a significantly (*p* = 0.0484) reduced ability to discriminate between the novel and familiar object (Figure 2. A). In the female cohort, both the saline-treated and MRG1061-treated mice could no longer discriminate between novel and familiar objects after 1 month (Fig 2. B). This model has previously been used to assess therapeutic potential in restoring NOR (**Greenfield et al**, **2022).**

**Figure 2.**
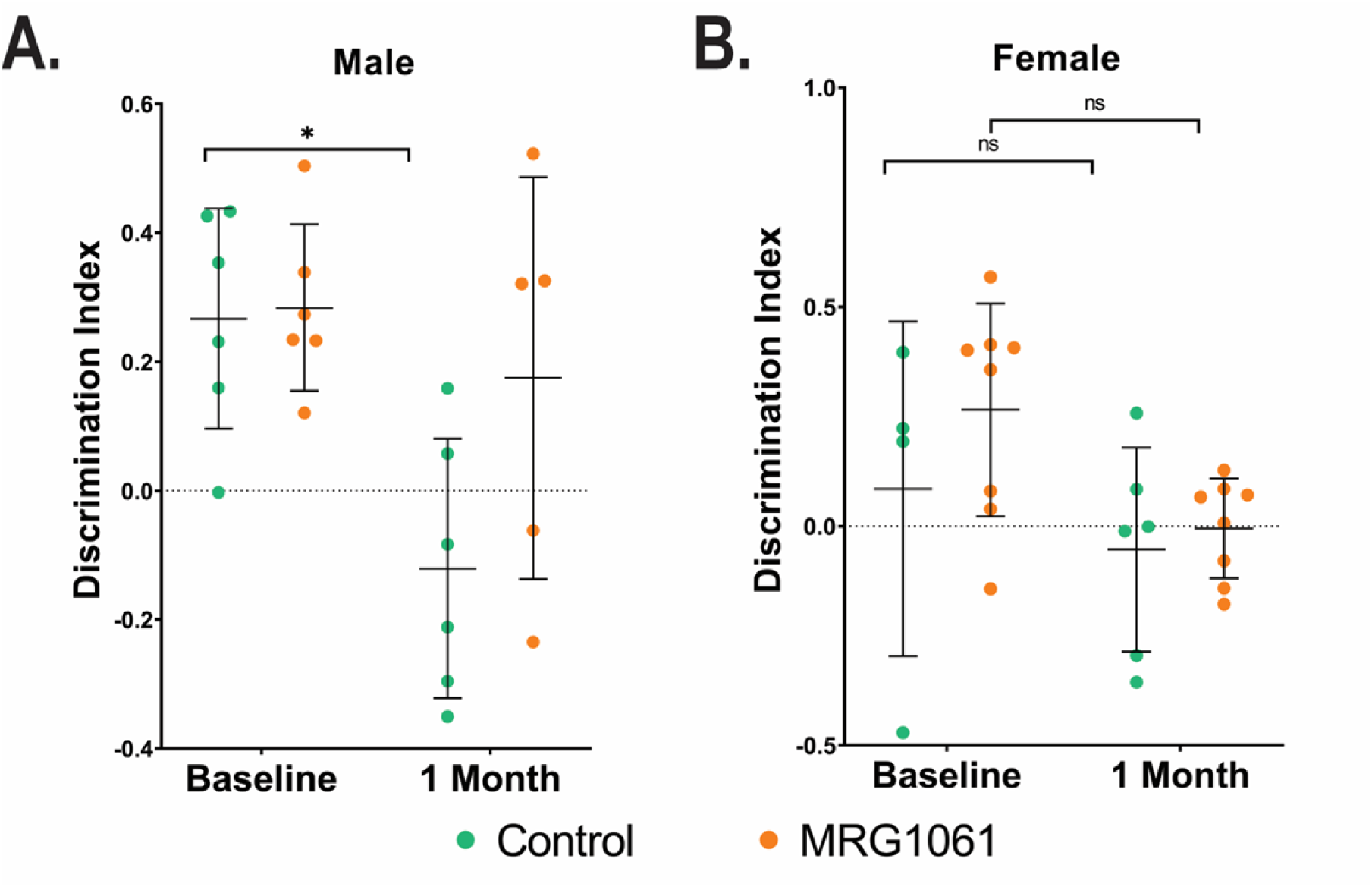
Novel object recognition. (A) Discrimination index at baseline and 1-month for male mice. *n*=12 Paired T-Test *< 0.05. (B) Discrimination index at baseline and 1-month in female mice. *n*=12

Efficacy of MRG1062 was tested in the motor neurone disease Q331K mouse model (**Mitchell et al., 2015).** Over the duration of the experiment, 3 animals did not survive the duration of the experiment, all 3 of the animals were in the control, saline treated animal group. All MRG1062-treated mice survived for the duration of the experimental protocol (data not shown). To test motor strength and locomotion, mice were tested on the Rotarod apparatus. In whole group analysis, both saline-treated Q331K mice and MRG1062-treated mice saw a significant decline (paired T-Test) over time in their ability to stay on the rotating platform as measured by change from baseline duration to fall. Saline-treated mice demonstrated a significant decline at 2 months from baseline in their ability remain on the apparatus (25.4% decline, baseline vs 2 months, p = 0.024) (Figure 3. A). In MRG1062-treated Q331K, there was also an age-related decline over two months in their ability to stay on the rotating platform (13.2% decline), this also reached significance (baseline versus 2 months) (Figure 3, A). For both groups there was no significant decline in ability between baseline and 1 month, with an accelerated decline between 1 month and 2 months. There was a 12.2% difference in decline from baseline in the saline treated group versus MRG1062-treated group at two months although this did not reach significance (unpaired T-Test). As we had determined a significant difference (unpaired T-Test, *p* = 0.0052), between male and female mice at baseline on the rotarod (data not shown) we analysed performance versus treatment by sex. There was no significant difference in reduction from baseline when comparing saline-treated male Q331K mice (23.5% reduction from baseline) with MRG1062-treated male mice (21.3% reduction from baseline), Figure 3. B. When we analysed female mice, there was a significant difference (*p* = 0.006) in reduction from baseline ability when comparing saline-treated mice at 2 months with MRG1062-treated mice at 2 months (Figure 3. C). Saline-treated female mice demonstrated a 26.8% reduction from baseline and MRG1062-treated mice demonstrated a 13.2% reduction from baseline.

**Figure 3.**
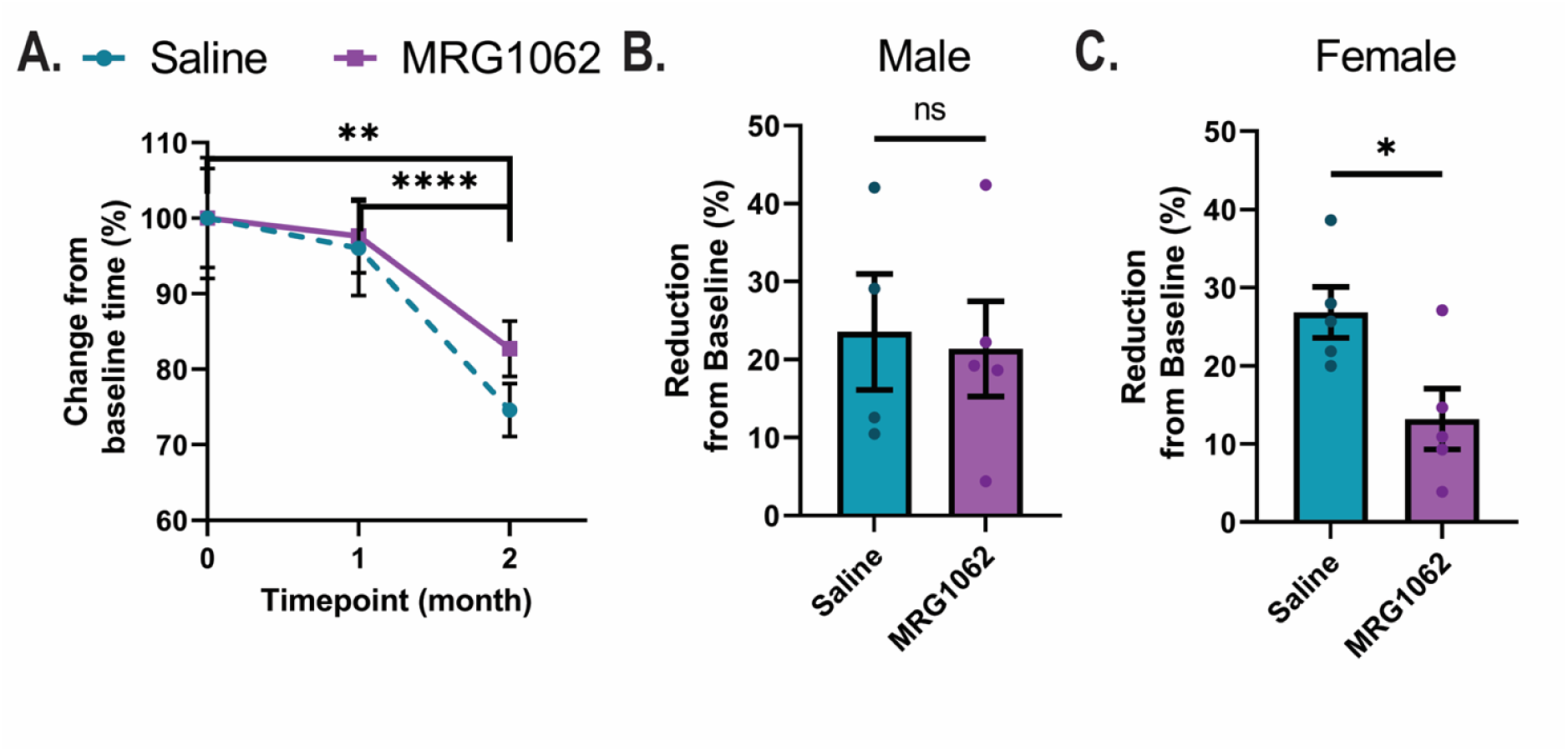
Effect of MRG1062 on motor function in TDP-43^Q331K^ mice. (A) Change in rotarod ability Q331K mice treated with MRG1062 vs saline over the course of the study. *n*=9 saline, *n*=10 MRG1062. Two-way ANOVA ** <0.005, **** <0.0001. (B) Quantitation of reduction from baseline in male saline and MRG1062-treated mice. *n*=4 saline, *n*=5 MRG1062. (C) Quantitation of reduction from baseline in female saline and MRG1062-treated mice. *n*=5 saline, *n*=5 MRG1062. Unpaired t-test *<0.05. Data are individual mice, error bars are SEM.

### 3.2 Effects on Pathology

There were no differences in brain weight between MRG1061-treated 5XFAD mice with saline-treated 5XFAD mice (data not shown). We examine the expression of full-length APP and low molecular species (LMW) of APP including Aβ. Figure 4 A, B show the quantitation of full-length APP and LMW APP. There was no significant difference (unpaired T-test) in any of the amyloid species, APP, LMW APP, and Aβ, when comparing MRG1061-treated mice with saline-treated animals. To examine regional expression in disease, immunohistochemistry of disease relevant areas was performed to examine APP and Aβ. The hippocampus, cortex, and pontine gray (pons) regions were selected. Figure 4. C shows representative images of immunohistochemical staining of Aβ in regions of interest demonstrating mice display pathological features consistent with Alzheimer’s disease. QuPath analysis to quantify staining of Aβ revealed no significant difference (unpaired T-test) in any Aβ staining when comparing MRG1061-treated mice with saline-treated animals (Figure 4. D, E, F).

**Figure 4.**
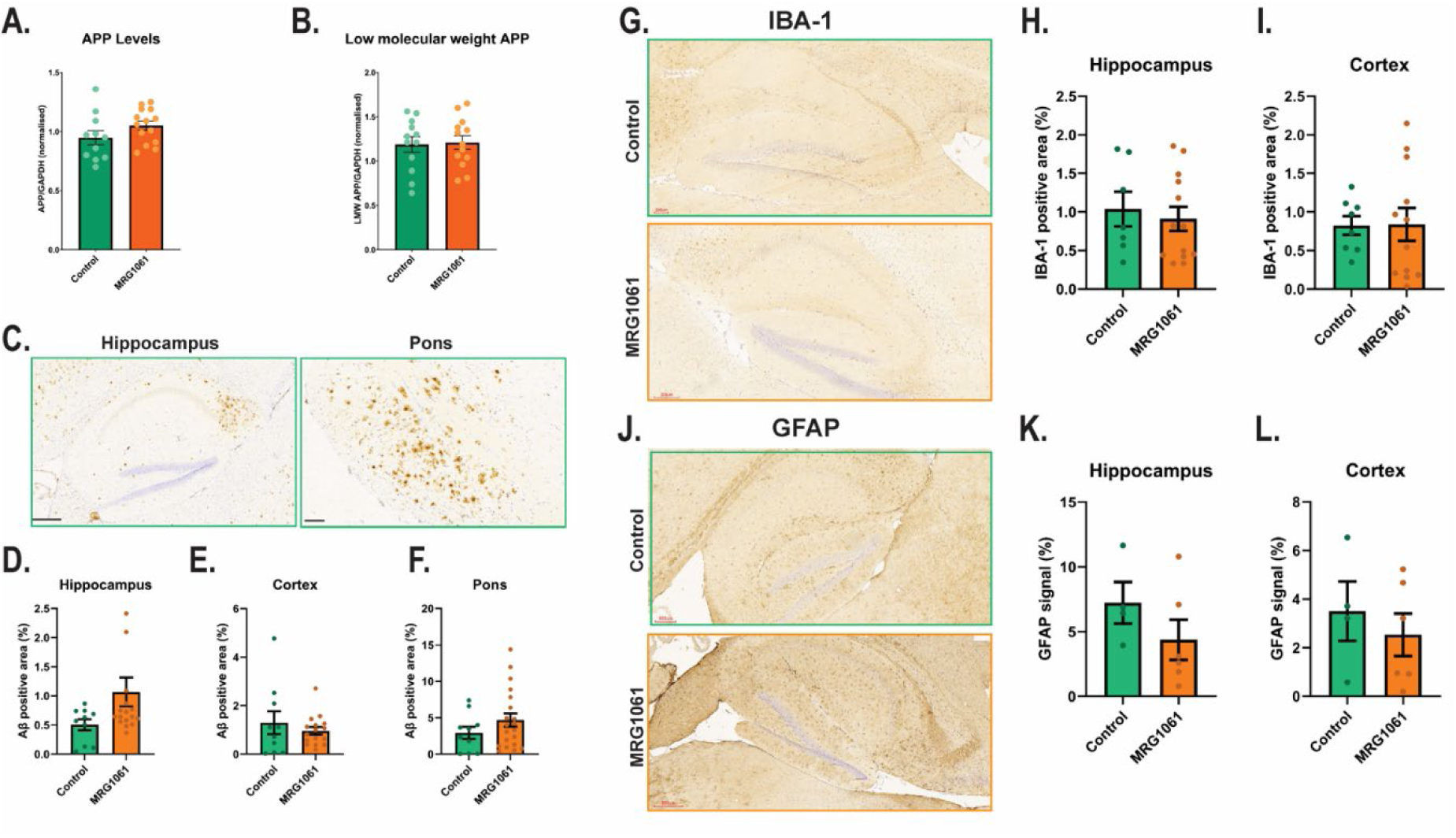
Effect of MRG1061 on amyloid levels in 5xFAD model. (A-B) Quantitation of full-length APP and LMW APP in 5XFAD mice. Total protein levels were normalised to GAPDH. Unpaired t-test. (C-F) Immunohistochemical analysis of 5XFAD mice. C. representative images of brain slices stained for Aβ. (D-F) QuPath quantitation of Aβ positivity across three different brain regions. Scale bar Hippocampus 300 µm, Pons 100 µm. Data are individual mice, error bars are SEM.

The major pathological feature of Motor Neurone Disease is the aggregation of the TDP-43 proteins. Western blotting of brain lysates from both treatment groups revealed no differences in TDP-43 expression levels. Further analysis of p62 (a protein involved in aggregation of TDP-43) expression levels also did not demonstrate any difference between groups. Finally, to investigate synaptic connectivity, we examined the expression level of the synaptic scaffolding post synaptic density 95 (PSD95) protein, again there were no differences between groups (data not shown). To determine regional expression in disease relevant areas, immunohistochemistry was performed to examine p62. The hippocampus, motor cortex, and pontine gray (pons) regions were selected. QuPath analysis to quantify staining of p62 revealed no significant difference (unpaired T-test) in staining in any of the brain regions examined when comparing MRG1062-treated mice with saline-treated animals (data not shown).

### 3.3 Effects on Inflammation

To explore possible alterations in inflammatory responses in the 5XFAD mouse model, brain slices were stained with the activated microglial marker IBA-1and the astrocytic marker GFAP (glial fibrillary acidic protein). Figure 4. G J, show representative staining of IBA-1 and GFAP respectively. There were no significant differences (unpaired T-test) in GFAP or IBA-1 staining when comparing MRG1061-treated mice with saline-treated animals in any of the brain regions (Figure 4. H, I, K, L).

Similar staining and quantitation in the MND mouse model revealed no significant difference in staining of IBA-1 in the cortex or pons region of the brain when comparing MRG1062-treated mice with saline-treated animals (Figure 5. B, C). There was no significant increase in GFAP staining in any brain region of MRG1062-treated mice compared to wildtype mice (Figure 5. D, E, F). There was a significant decrease in IBA-1 staining in the hippocampus of MRG1062-treated mice compared to saline-treated mice, one-way ANOVA with post-hoc analysis, *p* = 0.0279 (Figure 5. A).

**Figure 5.**
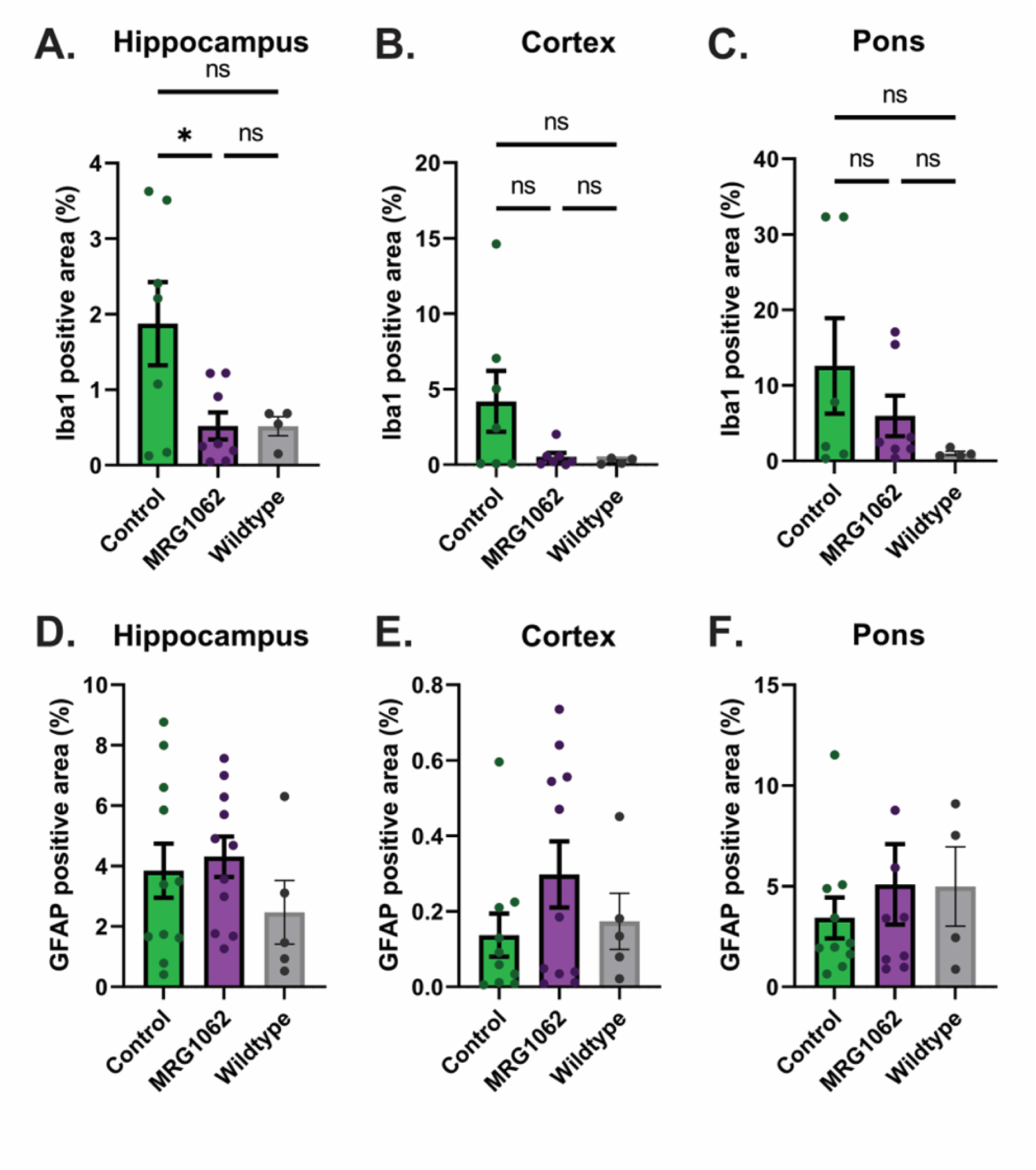
Quantification of IBA-1(A-C) and GFAP (D-F) staining of Q331K brain slices. Brain slices were stained with the microglial marker IBA-1 (A, B, C) or astrocytic marker GFAP (D, E, F) and % area positive determined for (A,D) hippocampus, (B, E) motor cortex and (C, F) pontine gray area. Statistics are One-way ANOVA with Tukey’s post-hoc.*<0.05. Data are individual mice, error bars are SEM.

### 3.4 Cytokine Analysis

As we determined alterations in the inflammatory response in the mouse model of motor neurone disease, we next investigated upstream mediators on inflammation in MRG1061 and MRG1062 treated mice brain tissue. The inflammatory profile was determined for both saline-treated and secretome-treated 5XFAD and Q331K mice brain lysates. Figure 6. B shows the heat map profile of the full cytokine panel tested for saline (control) and MRG1061-treated 5XFAD mice. We found a significant reduction in IL-1β in male MRG1061-treated mice compared to male saline-treated mice Figure 6. A (*p* = 0.0479).

**Figure 6.**
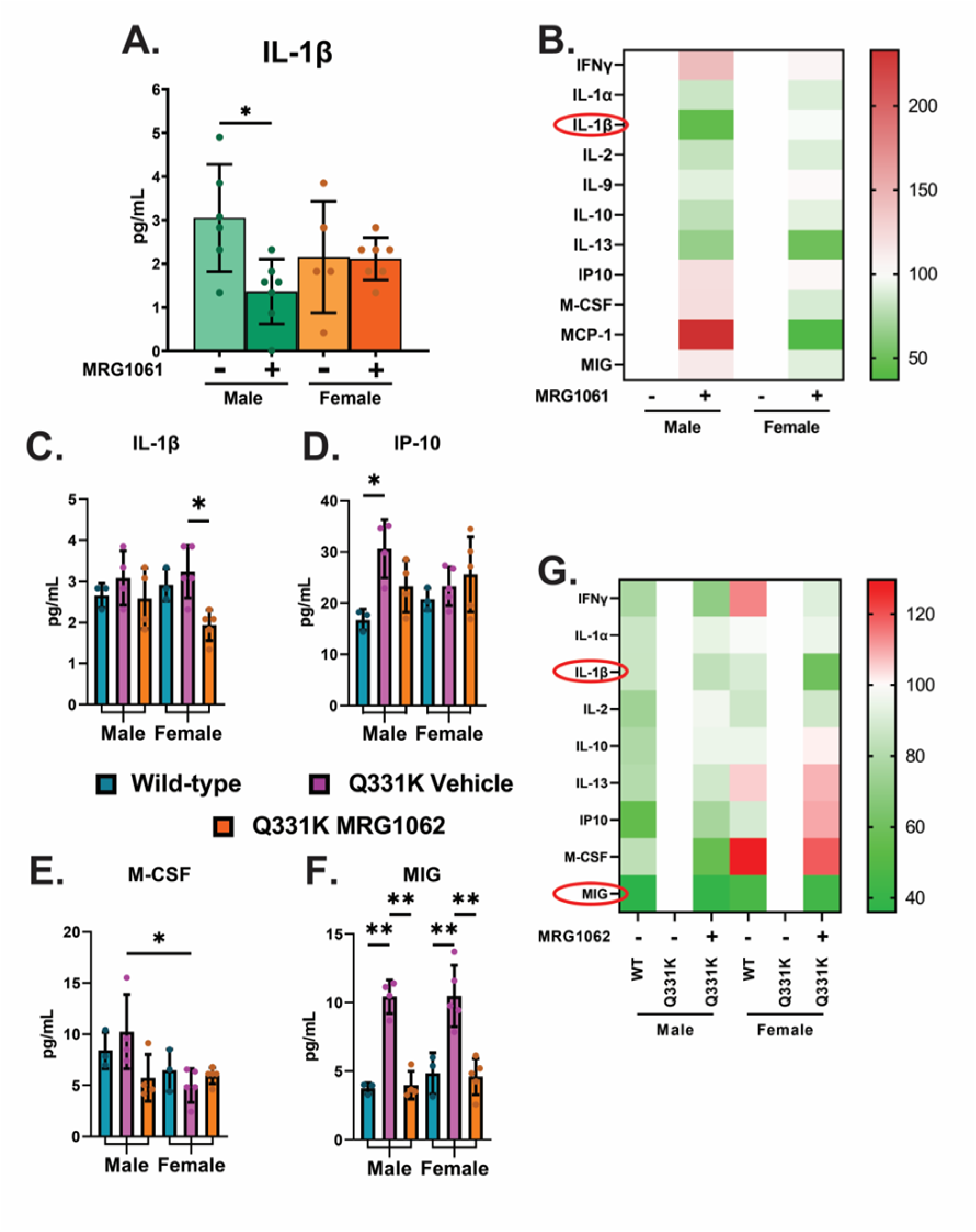
Protein expression levels of inflammatory cytokines within male and female brain tissue lysate following MRG1061/MRG1062 treatment. (A) Quantitation of IL-1β of cytokine protein concentration. (B) Heatmap of full multiplex panel. Data points are individual 5XFAD mice. *n*=5(male), *n*=6 (female) control, *n*=7 MRG1061 animals. Statistics are Two-way ANOVA with Tukey’s post-hoc. *<0.05. Bars are SEM. Quantification of inflammatory cytokine protein levels in brain tissue lysates from male and female Q331K mice following MRG1062 administration. (C, D, E, F). Individual graphs depict concentrations of selected cytokines. (J) Heatmap visualisation of multiplex cytokine panel. Data points are individual mice *n*=3 Wild-type, *n*=4 (male), n=5 (female) control, *n*=4 (male), *n*=5 (female) MRG1062. Statistics are Two-way ANOVA with Tukey’s post-hoc. *<0.05, **<0.01. Bars are SEM.

Similar to our behavioural analysis, cytokine profiling revealed sex differences in response to MRG1062 in the Q331K mice. Cytokine profiling revealed that MRG1062 significantly reduced IL-1β in female mice compared to saline-treated female mice (Figure 6. C, *p* = 0.0343) and significantly reduced MIG (CXCL9) in both male and female MRG1062-treated mice compared to saline treated (Figure 6 F, WT v saline male *p* = 0.0023, saline v MRG male *p* = 0.0012, WT v saline female *p* = 0.0049, saline v MRG female *p* = 0.0023). There were no significant differences between WT mice and MRG1062-treated male and female mice when measuring MIG (Figure 6. F).

### 3.5 Human Blood Testing Results

As we identified changes in cytokine and inflammatory markers in mouse models of neurodegeneration, we next investigated the translatability potential of MRG1061 and MRG1062 in human blood. Whole blood was preincubated with either MRG1061 or MRG1062 for 1 hour prior to stimulation. MRG1061 and MRG1062 significantly reduced the LPS-induced increases in Il-6 and TNFα (Figure 7. A, B. *p* < 0.0001) while significantly enhancing LPS mediated increases in MCP3, IL-10 and IL-1RA (Figure 7, E *p* <0.0005). Given the responses detected over LPS stimulated blood, we next investigated if MRG1061 and MRG1062 possessed an inherent anti-inflammatory action. Each secretome was incubated for 6 hours in whole blood and RPMI media (without LPS addition) and the cytokine profile determined. Secretome pretreatment significantly reduced IFNγ, *p* < 0.005 (Figure 7. F)

**Figure 7.**
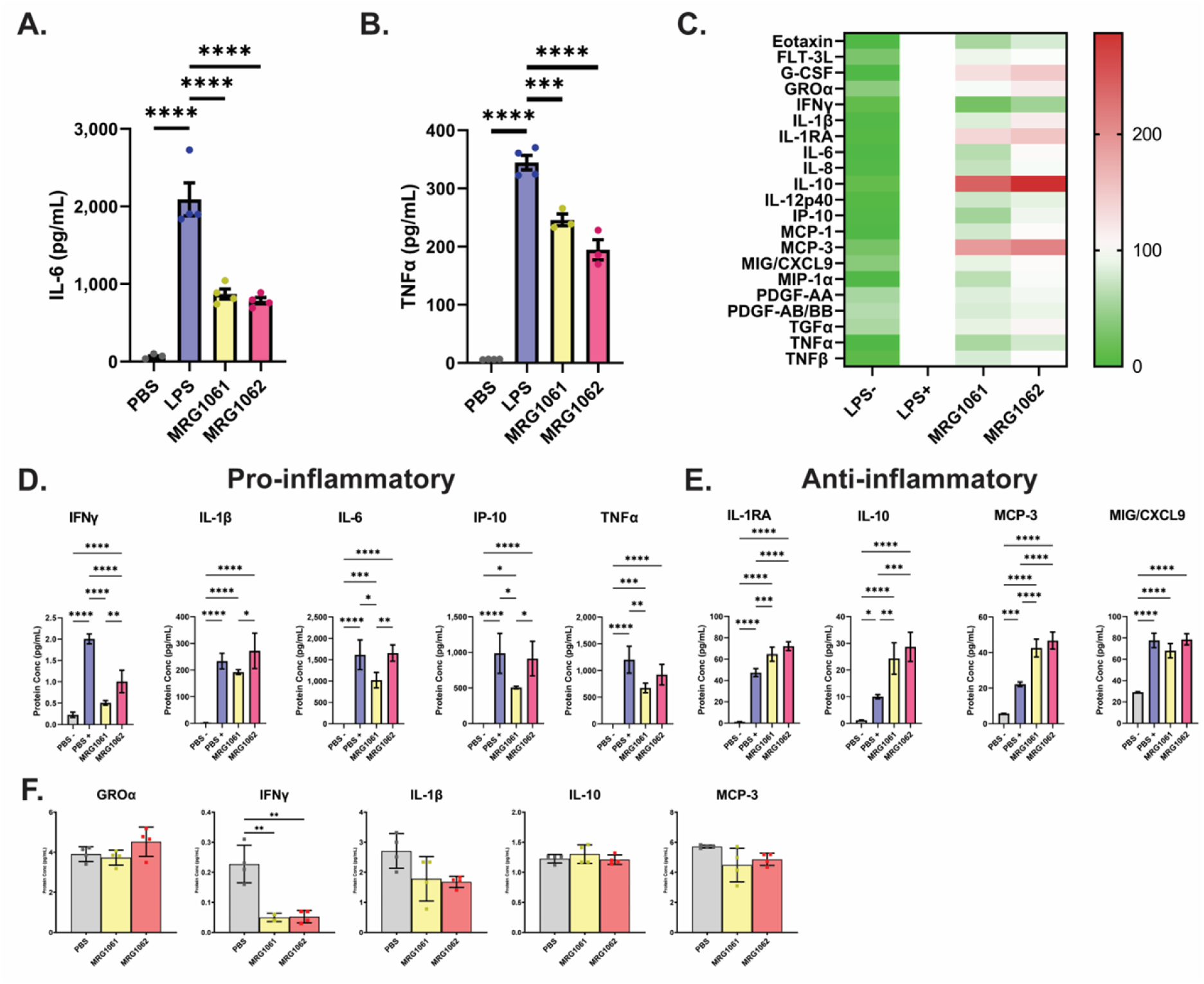
Ex vivo assessment of stimulated human inflammatory cytokine proteins from human blood following MRG1061 and MRG1062 administration. (A-B) Individual assessment of proinflammatory IL-6 and TNFα levels. (C) Visualisation of a broad multiplex cytokine/chemokine panel following ex vivo dosing of secretome formulations. (D) Individual graphs of pro-inflammatory proteins from multiplex assessment and (E) anti-inflammatory proteins. Data are isolated whole blood from 3 different human donors. Statistics are one-way ANOVA with Tukey’s post-hoc. *<0.05, **<0.01, ***<0.001, ****<0.0001. Bars are SEM. (F) Individual assessment of unstimulated circulating human inflammatory cytokine proteins following MRG1061 and MRG1062 administration. Data are isolated whole blood from 3 different human donors. Statistics are one-way ANOVA with Tukey’spost-hoc. **<0.01. Bars are SEM.

## 4. Discussion

Our current study demonstrates that exogenous application of a stem cell secretome can modulate behavioural phenotypes in two different models of neurodegenerative disease without effecting the classical neuropathological hallmarks. We first investigated the application of the stem cell secretome in an Alzheimer’s disease mouse model, the 5XFAD mouse (**Oakley et al, 2006**).

At baseline mice could discriminate between novel and familiar objects. When we analysed the NOR data at baseline and parsed by sex, female mice had a lower discrimination index ratio than male mice. After 1 month of saline treatment the male mice could no longer discriminate between novel objects and familiar objects suggesting that episodic memory was impaired, this loss of ability to discriminate was significant. The ability to discriminate was maintained in the male MRG1061-treated mice after 4 weeks and there was no significant difference between baseline and 1-month MRG1061 treatment in their ability to discriminate, suggesting that episodic dependent memory had been maintained in MRG1061-treated male mice. In the female cohort, there was no difference in the discrimination ability in the saline treated mice at baseline and after 1 month treatment and no difference in ability to discriminate between the saline-treated female mice and the MRG1061-treated female mice. Our observation on sex difference, at baseline in NOR, aligns with reported differences in pathology and behavioural defects in the 5XFAD mice being dependent on age and sex of the mice (**Padua et al, 2024; Sil et al, 2022**). In our study, mice were treated at 11 weeks age, where differences on spatial memory have previously been reported (**Oakley et al, 2006**). More recently it has been reported that female mice demonstrate an accelerated onset of pathology compared to male mice (**Poon et al, 2023**). As such female mice have a quicker onset of behavioural and pathological phenotype. It is possible that in our study the onset of pathology is accelerated in the female mice in comparison to male mice to a point where treatment with MRG1061 would no longer have any effect.

We next investigated motor impairment in Q331K mice on the rotarod. After 1 month of treatment with either saline or MRG1062 there was no significant impairment in performance on the rotarod as measured by change from baseline. After 2 months, both saline and MRG1062 treated mice displayed a significant decline in motor impairment compared to baseline and 1 month treatment. Although there was a greater decline in the saline treated mice compared to the secretome treated mice (25.5% vs 12.2% decline in ability) this did not reach significance. The impairment in the performance on the rotarod in the saline treated mice is similar to that reported previously by the creators of the mouse line (**Mitchell et al**, **2015**). Like Alzheimer’s disease mouse models, mouse models of motor neurone disease also display sex differences in disease progression and onset and severity of motor phenotypes (**Watkins et al, 2020**). When we analysed the motor performance by sex, we found that MRG-treated female mice had a significant reduction in rate of decline from baseline compared to saline-treated female mice. There was no difference in decline from baseline when comparing male saline-treated and male MRG1062-treated mice. It has previously been reported that female Q331K mice show less severe motor deficits than male mice (**Watkins et al, 2020**), while male mice display reduced rotarod performance around 6 months (**Watkins et al, 2021**). Similar to our results in the 5XFAD mice, the improved performance in female mice when compared to male mice may reflect a therapeutic window where pathology has not advanced to a point where interventions would have limited efficacy.

Given our observed differences in behavioural phenotypes in the MRG1061/62-treated animals we next examined underlying pathological markers in both the 5XFAD model (Aβ/APP) and the Q331K model (TDP-43) and determined no differences between animals treated with saline or MRG1061/62 respectively. There are limited studies for the effects of stem cell secretomes on autophagy and protein degradation specifically in the context of neurodegenerative diseases. MSC have been reported to enhance autophagy of Aβ in a neuronal cell model and an Alzheimer model with implanted MSCs (**Shin et al, 2014**) and chronic treatment of a C. elegans model of Parkinson’s disease offered protection against alpha-synuclein neurodegeneration (**Marques et al, 2021**). While there is an accumulation of reports that demonstrate how stem cell secretomes can impact on protein degradation, either through direct effects on proteostasis and the ubiquitin-proteosome systems, or by directly increasing autophagy (**Daneshmandi et a, 2020**), there is no direct evidence to support increased clearance of pathological hallmarks of Alzheimer’s disease (amyloid plaques or tau tangles) or Motor Neurone Disease (TDP-43 accumulation). There is however a growing body of work to support stem cell secretomes in modulating inflammatory responses. GFAP has been used as a biomarker of neuroinflammation in multiple neurodegenerative models to indicate astrocyte activity. Elevated levels have been reported in 5XFAD mice (**Manji et al, 2019**) however in our study we found no difference in GFAP staining between saline-treated 5XFAD mice and MRG1061-treated mice. IBA-1, a marker of activated microglia, has been reported to be increased in 5XFAD mice compared to WT mice at 10 months of age (**Yoo et al, 2023**). Although other reports find that IBA-1 positive microglia are associated with AB deposits as early as 10 weeks (**Boza-Serrano et al, 2018**). In our study we did not detect any change in IBA-1 staining in either treatment groups in the 5XFAD mice.

When we examined the brains from saline or MRG1062-treated Q331K mice and untreated wild type (WT) mice littermates, we found no differences in GFAP staining in any brain region examined. These findings are not unexpected as other studies have also reported no change in GFAP staining across multiple brain regions in Q331K mice (**Lin et al, 2021**). When we examined the brains from WT and Q331K mice we found a trend towards increased IBA-1 staining in the saline treated-Q331K compared to untreated WT mice in all brain regions examined and reduced IBA-1 staining in the MRG1062-treated Q331K mice compared to saline-treated Q331K mice. In all brain regions, the IBA-1 staining was reduced to an amount comparable to untreated WT. While there was a clear reduction in IBA-1 staining in MRG1062-treated Q331K mice compared to saline-treated Q331K mice in all brain regions examined, this difference only met significance in the hippocampus. This suggests that in the Q331K model of motor neurone disease, glial involvement is predominantly microglial as previously reported (**Lin et al, 2012**) and that MRG1062 can moderate microglial activation in this model.

We next examined mediators of inflammation in our mouse models. Cytokine analysis of brain lysates revealed that in the 5XFAD mice there was a sex dependent significant reduction in IL-1β in the MRG1061 treated male mice compared to saline-treated male mice. There were no differences detected in the female mice. IL-1β has been reported to mediate AD pathogenesis through a number of different mechanisms, such as increased pathology (**Ghosh et al, 2013**) and Aβ clearance through upregulation of genes involved in immune response (**Rivera-Escalera et al, 2010**). Of note, sex specific effect of MRG1061 on cytokine levels and behavioural tests was evident only in the male mice. When we examined the same panel of cytokines in the Q331K mice we found a similar sex dependent effect of MRG1062 on IL-1β levels. Specifically, there was a significant reduction in IL-1β in MRG1062-treated female mice compared to saline treated female mice. There was no effect of treatment in the male mice. Again, the modulatory effect of MRG1062 on IL-1β in female mice mirrored the modulatory effect of MRG1062 on motor coordination in that the effect was specific to the female cohort. We also found a significant increase in IP-10 in the male saline-treated Q331K mice compared to WT untreated mice. Treatment with MRG1062 reduced the levels of IP-10 in male mice although this did not reach significance. IP-10 has been used a biomarker for motor neurone disease, and its upregulation is linked to disease progression in other neurological conditions (**Wang et al, 2018**). The increase in IP-10 would support our observation that in the male cohort of Q331K mice that the disease had progressed more than the female mice reducing the therapeutic potential of MRG1062 in advanced stages of the disease. Supporting this, macrophage colony-stimulating factor (M-CSF) has been linked to motor neurone disease and other neurodegenerative conditions as a marker of neuronal damage (**Barbieto et al, 2010**). In our Q331K mice, there was a significant upregulation of M-CSF in the male mice compared to female mice. The inflammatory cytokine monokine induced by interferon-gamma (MIG) was significantly elevated in both male and female Q331K mice compared to WT untreated mice and treatment with MRG1062 significantly reduced MIG in both male and female mice. MIG is an inflammatory cytokine induced by interferon gamma (IFNγ) (**Mahalingam et al, 2000**). IFNγ can induce pathological hallmarks of motor neurone disease in cell models (**Chun et al, 2024**) and elevated levels have been reported in serum and CSF samples from motor neurone disease patients (**Liu et al, 2024**).

Given the effect of MRG1061/1062 on inflammation and its mediators in our animal models we investigated the potential of each secretome to modulate pro and anti-inflammatory mediators in whole human blood stimulated with lipopolysaccharide (LPS). In general, MRG1061 and to a smaller degree MRG1062 significantly reduced proinflammatory mediators IFNγ, IL-6, IP-10 and TNFα, and significantly induced anti-inflammatory mediators IL-1RA, IL-10 and MCP-3.

In both models, MRG1061/62 reduced IL-1β and microglial activation concomitant with ameliorating behavioural phenotypes. These effects were sex-dependent in both animals and possibly linked to extent of underlying pathology and suggest a therapeutic window in both models for improved efficacy. MRG1061 and MRG1062 were able to ameliorate these effects without impacting on underlying pathological drivers. A number of studies have reported that different approaches to treating mouse models of neurodegenerative conditions can impact on behavioural phenotypes without addressing the underlying pathology. This has been achieved through a number of strategies, such as environmental enrichment (**Jankoswky et al, 2005**), enhancing synaptic plasticity (**Pham and Dore, 2023**) or directly targeting the sequalae of following the onset of pathology, such as inflammation (**Spangenberg et al, 2016**).

To our knowledge, this is the first report of a novel therapeutic candidate ameliorating both biochemical and behavioural phenotypes in two different models of neurodegeneration without impacting on pathological hallmarks. It is now accepted that underlying pathology can occur up to 30 years before clinical phenotypes present (**Johnson et al, 2023**).

Following our findings, observing an ability for stem cell secretomes to modulate behavioural deficit and inflammatory responses in neurodegenerative murine models without directly altering standard neuropathological markers, this preliminary study supports a strategy for a use as a potential preventative strategy. The MSC secretome’s multifaceted actions, including reducing neuroinflammation and promoting neuroprotection, offer a novel approach to delay or mitigate disease progression before irreversible pathology develops. This is especially relevant considering that neurodegenerative pathology such as amyloid plaques and protein aggregates can precede clinical symptoms by decades. Stem cell secretome-based therapies may therefore provide a non-invasive, cell-free intervention to preserve cognitive and motor functions in at-risk populations prior to onset of detectable disease, potentially extending the therapeutic window and improving patient outcome.

## Acknowledgements

The authors would like to thank Jackie Mitchell, Ph.D, King’s College London for the Q331K mice and her advice on related motor experiments. We would like to thank Paul Armstrong, Ph.D, University of Leeds for their guidance and training on the rotarod experiments. Finally, we would like to thank Barry Sharples (Micregen Ltd.) for their careful review of the manuscript and acquisition of funding. Funding for the work was proved by a Medical Research Council Confidence in Concept award to the University of Bradford and matched funding by Micregen Ltd.

## CRediT authorship statement

**Stuart Dickens**: Investigation, Formal Analysis, Methodology, Project administration, Visualization, Writing – review and editing. **Andrew Parnell**: Formal Analysis, Investigation, Methodology, Writing – reviewing and editing. **Daniel Feist**: Formal Analysis, Investigation, Methodology. **Ben Mellows**: Formal Analysis, Methodology, Writing – reviewing and editing. **Ketan Patel**: Conceptualization, Supervision, Writing – reviewing and editing. **Steve Ray:** Conceptualization, Supervision, Writing – reviewing and editing. **Samantha McLean**: Conceptualization, Supervision, Funding acquisition, Writing – reviewing and editing. **Robert Mitchell**: Formal Analysis, Investigation, Methodology, Supervision, Visualization, validation, Writing – reviewing and editing. **Ritchie Williamson**: Conceptualization, Investigation, Project administration, Supervision, Funding acquisition, Formal analysis, Writing – Original draft

## Nonstandard abbreviations

Aβ: Amyloid beta
AFSC: amniotic fluid-derived mesenchymal stem cells
APP: amyloid precursor protein
AD: Alzheimer’s disease
GFAP: Glial fibrillary acidic protein
MCSF: macrophage colony-stimulating factor
MSC: multipotent stromal cells
MND: Motor Neurone Disease
NOR: novel object recognition
PSD95: post synaptic density 95
Recommended section: Drug Discovery and Translational Medicine

